# Myocyte Damage and Mass Loss Drives Increasing Water Content, Compliance, and Survival in Septic Cardiomyopathy

**DOI:** 10.1101/2025.07.07.663608

**Authors:** Verity J Ford, Steven B. Solomon, Junfeng Sun, Willard N Applefeld, Jing Feng, Jeffrey Wang, Irene Cortes-Puch, Yan Li, Michael A Solomon, Zaid N. Safiullah, Robert L Danner, Marcus Y Chen, Parizad Torabi-Parizi, Harvey G. Klein, Zu-Xi Yu, Charles Natanson

**Author notes:** **Corresponding Author:** Charles Natanson, MD, Critical Care Medicine Department, NIH Clinical Center, 10 Center Drive, Room 2C145, Bethesda, MD 20892. These two should be equally considered first authors (Verity J Ford, Steven B. Solomon). **Conflicts of interest:** The authors have declared that no conflict of interest exists. The work by the authors was done as a part of the US government-funded research; however, the opinions expressed are not necessarily those of the NIH.

## Abstract

**Introduction/Purpose:** During the septic cardiomyopathy, the mechanism and relationship to outcome of changes in left ventricular (LV) end diastolic volume (EDV) and ejection fraction (EF) remains obscure. We compared serial changes in LVEF and LVEDV to successive alterations in LV wall ultrastructure, water content, and total mass to investigate whether these measures can explain their basis.

**Methods:** We performed cardiac magnetic resonance imaging at 0,6,18,30,42,54, and 92h post-bacterial challenge in a large-animal model (n=57) that mimics human septic cardiomyopathy. LV tissue was obtained for electron microscopy (EM) upon death and 66h in sacrificed survivors.

**Results:** Between 0-6h post-challenge, LV compliance and EDV reached its greatest decline. Non-survivors (n=18) exhibited significantly greater reductions in LVEDV, along with more myocyte edema, mitochondrial swelling and myofilament fragmentation on EM. This increased tissue damage may explain why non-survivors developed worse LV compliance and a greater decline in LVEDV, which persisted until death. From 6-30h, LVEDV significantly improved to baseline in non-survivors, while survivors experienced ∼20% increases (n=39). Concurrently, there was significant LV mass loss and increases in percent water content that were significantly associated with increases in LVEDV. This is consistent with a passive mechanism for rapidly improving LV compliance and EDV. Full recovery of EF required additional days. We hypothesize the prolonged significant mass loss over 5d reflects an active process for remodeling fragmented myofilaments, eliminating myocyte edema, and mitochondrial swelling, ultimately restoring contractile function.

**Conclusion:** The septic cardiomyopathy constitutes a diffuse ultrastructural injury to myocytes with three phases. Initially, there is a decrease in LVEDV, and EF due to myocyte damage within 6h of bacterial challenge; next, the patient sees a passive LVEDV recovery from 6-30h, where LV mass loss increases relative wall percent water content, which facilitates wall compliance and LVEDV; and lastly, the patient sees mass loss beyond 30h consistent with an active repair mechanism of myocytes, returning systolic function to normal. Therefore, EDV changes are a pathophysiological biomarker for sepsis outcomes. A lower LVEDV indicates persistent unrepairable ultrastructure damage with worsening wall compliance and poorer outcomes. LVEDV dilation is a sign of near-full recovery of ultrastructure injury, augmenting wall compliance and improving outcomes.

**Clinical Implications:** We explain herein why septic cardiomyopathy findings don’t have clinical implications like heart failure. Septic patients who exhibit signs of heart failure, low LVEF with high EDVs, are doing well – reflecting mild myocyte injury, effective damaged tissues clearance, increased relative LV wall water content, and compliance. This augments the LVEDV, lowering the LVEF. Septic patients who deteriorate rapidly, contrary to heart failure patients, show high/normal LVEF and low/normal LVEDV. Here, the myocyte damage is severe, leading to insufficient wall repair, and this decreased wall compliance persists, preventing the LV from dilating and making LVEDV low which ultimately raises the LVEF.

## Introduction

The “Septic cardiomyopathy” was first characterized at the National Institutes of Health (NIH) in the 1980s in both humans and a large animal model.^1,2^ Researchers employed daily Cineangiogram, which used radiolabeled red blood cells and gamma cameras gated to heart rate to calculate the left ventricular ejection fraction (LVEF) and thermodilution Swan-Ganz catheters to quantify stroke volume (SV), collectively used to determine a calculated end diastolic volume (EDV)(i.e., (EF /SV =EDV).^1,2^ A prominent feature of this condition is a transient reduction in LVEF, which can fall from a normal range of 50–65% to as low as 25– 30% within 2-3 days following the onset of septic shock.^1,2^ In survivors, this period is marked by ventricular dilation (increase in LVEDV), with both EF and LVEDV typically returning to pre-sepsis levels over 10-14 days.^1,2^ However, the underlying mechanisms leading to this cardiac dysfunction—and the reasons why some patients reversibly dilate their ventricles causing the LVEDV to increase and survive, while others do not and die—remains poorly understood, with previous studies proposing a variety of unconfirmed hypotheses.^3^

Using a large-animal model that closely mirrors human septic cardiomyopathy,^2,4,5^ we recently published studies using cardiac magnetic resonance images (cMRI) at baseline, 48h, and 96h after bacterial challenge.^6,7^ Daily echocardiograms, intracardiac pressure measurements, and LV electron micrographs (EM; sampled at 66h) complemented the imaging data.^7^ We found that at 24h post-challenge, echocardiography revealed a significant profound reduction in ventricular chamber size in non-survivors that was significantly greater than what was observed in both survivors and controls. These LVEDV findings could not be explained by differences in preload, afterload or heart rate changes. During the subsequent 24h—the period of maximal EF decline— both survivors and non-survivors developed increases in ventricular chamber size. By 48h, cMRI demonstrated a pronounced increase in LV wall edema (cMRI T2), which light microscopy and EM at 66h confirmed as the dominant histopathological feature. Consistent with these experimental observations, myocardial edema has also been documented in patients during septic shock, underscoring a potential role of edema in septic cardiomyopathy.^3,8,9^ Moreover, myocardial edema has been associated with poor outcomes in other cardiac pathologies, such as myocardial infarction and Fabry disease, further supporting its role as a marker of injury severity and impaired myocardial function.^10,11^

From this, we hypothesize that myocardial edema may be a key factor in early cardiac dysfunction in sepsis and/or later recovery. To test this hypothesis, we conducted a detailed time-course study by collecting serial cMRI at baseline, 6,18,30,42,54, and 92h following a lethal bacterial challenge. Post-mortem LV tissue was also collected for EM and light microscopy. We sought to identify, through cMRI and histology, whether there is a temporal relationship between outcome, the development of myocardial edema, and the early decreases and/or later increases in LVEDV or the falls in LVEF.

## Materials and Methods

### Study Design and Animal Model

We utilized a well-established canine model of bacterial pneumonia to assess the effects of sepsis on cardiac function comparing survivors and non-survivors^2,4,5^ ( see Figure S1 for study timeline). Purpose-bred beagles (9-15 kg, 18-30 months old, male, Marshall Farms, Concord, MA) were sedated, tracheostomized, and mechanically ventilated on day one, and then at 0h, animals received an intrabronchial challenge of *Staphylococcus aureus* (0.6–1.2×10⁹ CFU/kg) to induce sepsis as previously described.^2,4,5^ All animals received comprehensive intensive care for up to 5d (survivors) not unlike titrated care as done around the clock clinically in human and large animal veterinary intensive care units (ICU).^4,5^ Briefly, animals were continuously monitored by trained technicians or physicians and sedation and analgesia was titrated to specific endpoints as previously described.^4,5^ Hemodynamic parameters were continuously monitored using percutaneously placed femoral arterial and pulmonary artery catheters including mean arterial pressure, central venous pressure, pulmonary artery pressure, pulmonary artery occlusion pressure, cardiac output, and heart rate. Standard care included continuous maintenance fluids (2 ml/kg/hour of Normasol-M with 5% Dextrose, ICU Medical. Lake Forest) supplemented with potassium chloride (27mEq/l, Fresenivs-Kabi, Lake Zurich, IL), given for five days or until death. To keep the pulmonary artery occlusion pressure above 10 mmHg, boluses of fluids were given every 2h as needed (20 ml/kg, NaCl, ICU Medical, San Clemente, CA) to maintain intravascular full resuscitation.^4–7^ The fractional inspired oxygen concentration, tidal volume, positive end expired pressure and number of breaths given were titrated to specific end points based on blood gas measurements that were obtained every 12h.^4–7^ Animal position was changed every 12h and intravenous famotidine (1mg/kg, q12, Fresenivs-Kabi, Lake Zurich) and subcutaneous heparin (3000IU, q8, Hospira, Inc., Lake Forest, IL) were given to prevent stasis ulcers and venous stasis clots, respectively. Intravenous ceftriaxone (50 mg/kg, Hospira, Inc., Lake Forest, IL), a *Staphylococcus aureus* sensitive antibiotic, was administered daily starting 4h after bacterial challenge. To delineate the underlying mechanism of sepsis-induced cardiomyopathy without the confounding effects of exogenous catecholamines on cardiac function, catecholamines were not used. Two animals were studied each week for up to 92 h (5 days) whereupon those alive would be considered survivors. A total of 57 animals were used across two studies. In the first study, the animals were treated identical to the second study except in the first study (n=29), animals received fewer cMRI then in the second study (n=28) (three vs. seven scans). The animals in the first study except for EM data in non-survivors were previously published.^6,7^ A subset of animals was euthanized (n= 7) at 66 h. These animals and survivors to 92h were euthanized (Euthaphen 75mg/kg, Sodium Pentobarbital (390mg/ml) and phenytoin sodium (50 mg/ml), IV, Dechra, Overland Park, KS). In all animals immediately after confirming death, tissue was obtained for histology and EM analysis.

See Supplement for expanded methods and study limitations.

## Results

### In the first 6h of septic shock, myocardial compliance maximally decreases more so in non-survivors: a severe injury phase

Within 6h after bacterial challenge, non-survivors exhibited the most severe impairment in LV relaxation and distention, as evidenced by the greatest decline in mean LVEDV (Figure 1A). This early decline at 6h is significantly more severe in non-survivors than in survivors and the mean LVEDV remains similarly significantly lower until death. The persistent decline at 6h, which remains until death, suggests a severe, injurious mechanism occurs during this acute period (0-6h). The recovery of LV wall distention and accommodation of filling toward more normal or “supernormal” EDVs begins after 6h. The rate of increases in LV EDV from 6-30h are similar and in parallel in survivors and non-survivors. This change likely reflects an outcome-independent recovery mechanism. However, a markedly less severe EDV decline at 6h being the case in survivors, with a similar rate of rise in EDV from 6-30h, survivors mean LVEDV not only return to near baseline as seen in non-survivors but exceed it by approximately 20%, and this elevation persists through to 96h. Reflecting in survivors a greater LV compliance overall or “super” recovery. The end systolic volume in the LV, mirrors the changes in EDV in survivors and non-survivors alike (Figure 1B).

**Figure 1.**
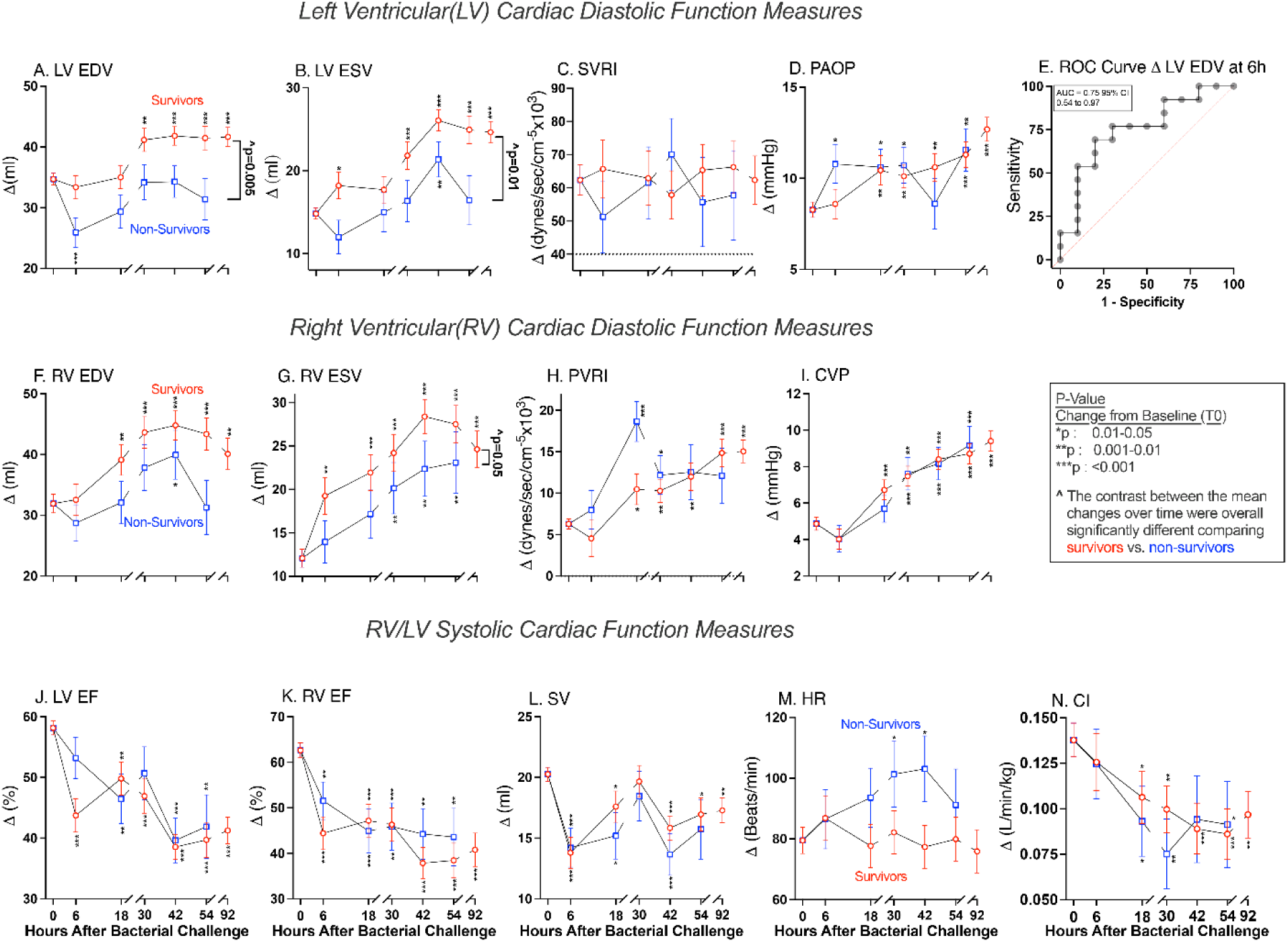
Diastolic and Systolic Function in Septic Shock. Serial mean (+/-SE) changes from baseline in cardiac and hemodynamic parameters are plotted from a common origin (mean baseline value for all animals). In Panels A-I shows cMRI and hemodynamic mean changes in diastolic function in large animal septic survivors (open red circles) and non-survivors (open blue squares). The serial means values for the left ventricle diastolic functions are shown in the top panels (Panels A-D). In Panel E the change in EDV from baseline to six h ability to predict survival is shown. The mean serial values for diastolic function of the right ventricle are shown on the middle panels (Panels F-I). In Panels J-N bottom row are shown different serial mean changes in cardiac parameters representing right and left ventricle systolic function. EDV, end diastolic function ESV, end systolic function SVRI, systemic vascular resistance index PAOP, pulmonary artery occlusion pressure ROC, receiver operating characteristic AUC, area under the curve CI, Cardiac Index PVRI, pulmonary vascular resistance index CVP, central venous pressure EF, ejection fraction SV, stroke volume HR, heart rate

As shown previously,^7^ differences in afterload (Figure 1C) and preload (Figure 1D) cannot easily explain the LV chamber size decline isolating our EDV decreases over 6h to a severe sepsis-induced compliance abnormality of the LV wall. Notably, these decreases in compliance by 6h predicted death (Figure 1E). Further the EDV changes over time in the RV virtually mirrored changes in the LV and these differences in EDV in non-survivors from survivors persist until death (Figure 1F-I). Taken together, these data indicate that early decreases in EDV are a critical initial damaging 6h long injury phase of the global sepsis induced cardiomyopathy.

### Systolic Recovery Lags Behind Diastolic Recovery

By 6h, survivors EFs were significantly lower in both ventricles (Figure 1J&K) with the nadir at 42-54h. Notably, prior studies in human septic shock suggest that, paradoxically, in some non-survivors, the LVEF may not significantly fall,^1,12,13^ we found this to be true in septic animals at 6h (Figure 1J). The more severe abnormality in wall compliance early on in non-survivors lowering LV and RVEDV (Figure 1A&F) appears to cause this lack of fall in RV & LVEF (Figure 1J&K). The less compliant wall prevents the ventricle from distending and filling in non-survivors compared to survivors. However, while SV is significantly depressed similarly in survivors and non-survivors at 6h (Figure 1L), the EF appears inexplicably normal in non-survivors. This is because the smaller EDV in non-survivors increases the SV portion of the EF ratio such that EF and performance are falsely elevated. The small number of reported cases of falsely elevated EFs in non-survivors may be because it is a time-related event and difficult to capture as it only occurs early on.

After 6h, the EF remains comparable between non-survivors and survivors throughout (Figure 1J&K), whereas the EDV is significantly and consistently lower in non-survivors throughout (Figure 1A). This finding can be attributed to a proportionally smaller SV and EDV in non-survivors, causing the EF ratio to remain comparable to survivors, who exhibit a larger SV and a larger EDV but similar ratio of the two parameters.

The last parameter that is independent of the changes in compliance of the LV wall that could possibly explain the alterations in LVEDV and EF is Heart Rate (HR). However, consistent with our previous studies,^7^ the mean HRs are not significantly different (∼80-110 beats/minute) between survivors and non-survivors throughout — and are not in a range fast enough to prevent cardiac filling and lower EDV or elevate the EF (Figure 1M).^14^ However, the mean HR was significantly increased from baseline at 30-42h in non-survivors. Overall, this higher HR suggests that the inability of non-survivors to augment SV by ventricular dilatation makes them more dependent on HR as a compensatory mechanism for a falling EF to maintain cardiac performance (Cardiac Index) during sepsis (Figure 1N).

### Increases in EDV, Not EF, is Associated with Survival

Given that in our animal sepsis model reductions in LVEDV by 6h predict death, whereas increases in LVEDV above pre-sepsis levels correlate with survival—and most of the serial changes in LVEF are not linked to outcome—we sought to determine whether these observations were clinically relevant. We previously published a meta-analysis examining 21 human sepsis cardiac studies published from 1983-2024.^7^ Here, we conducted an additional analysis of these data using weighted linear regression to assess if there are any differences in any of the studies between survivors and non-survivors over the full range of reported EFs and EDVs. Regardless of whether the mean EF of one of the 21 studies was low∼30% or high ∼60%, the regression line and its 95% confidence interval consistently encompassed every one of the 21 studies mean EFs, indicating that mean EFs did not differ between survivors and non-survivors over the broad observed range in 21 independent cardiac sepsis studies (Figure 2A). Eleven of these cardiac sepsis studies provided analyzable mean EDV and outcomes.^7^ Over the full range of mean EDVs observed across the 11 studies, the regression line and its 95 % confidence interval bounds were below the identity line, indicating that mean EDVs were lower in non-survivors compared to survivors (Figure 2B). Like our sepsis model, LVEF changes in clinical studies were comparable in survivors and non-survivors while EDVs were significantly higher in survivors and lower in non-survivors.

**Figure 2.**
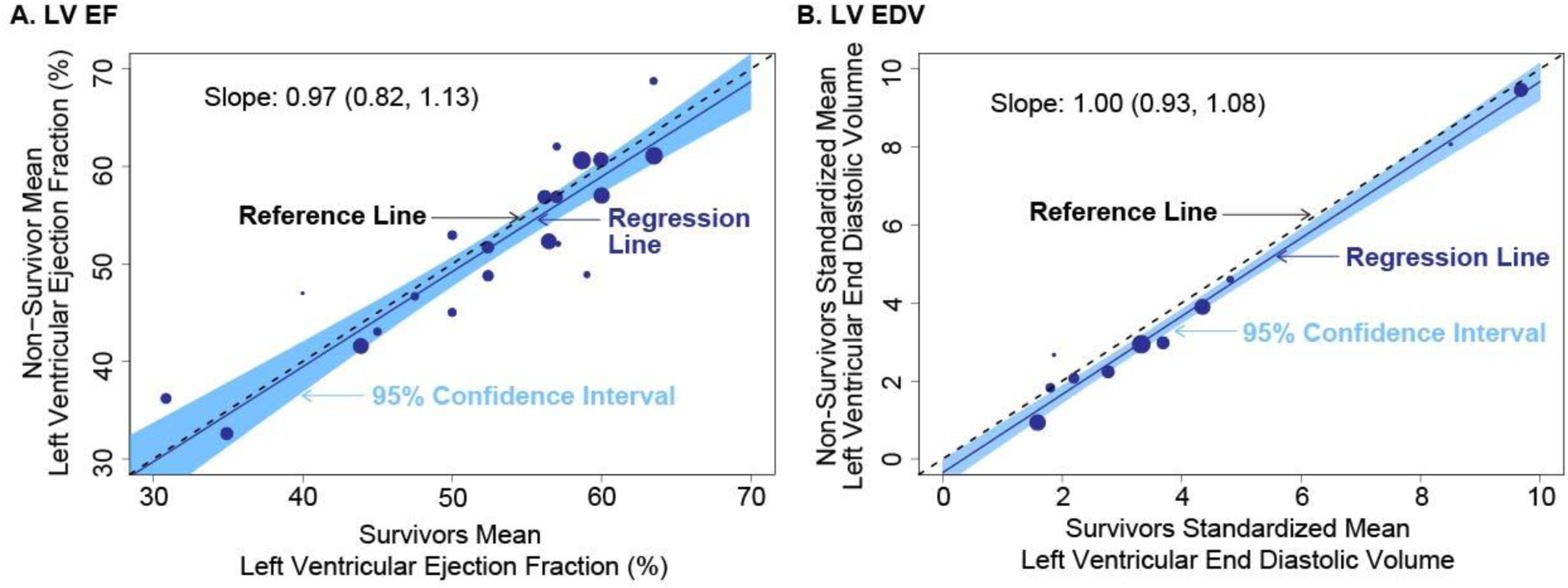
Cardiac LVEF and EDV in Septic Humans. Panel A shows mean cardiac function values from a published metanalysis of 21 cardiac studies in humans with septic shock from 1983 to 2024.^7^ Here a weighted linear regression analysis compared the mean LVEF in septic survivors (x-axis) verses septic non-survivors (y-axis) over the full ranges seen in these studies. The area of each circle is proportional to its weight (i.e., inverse variance) in the model. The 95% confidence interval bounds (blue region) for the regression line (blue solid line) for the relationship between survivors and non-survivors completely overlaps the identity line (dashed black line). This indicates in septic survivors regardless of the mean values for ejection fraction they were similar in septic non-survivors over all 21 studies. Eleven of these studies were evaluable and reported mean values for LV EDV (Panel B). The 95% confidence interval bounds fall below the identity line indicating regardless of the mean EDV in septic survivors, the non-survivors overall had a significantly smaller mean EDV across these 11 studies.

These experimental animal and clinical data taken together show that, in septic shock, the LVEDV is a potentially powerful useful biomarker to gauge outcome and is much superior to the LVEF. These data indicate that if the LV chamber dilates and the EDV increases out of the normal ranges in septic patients, survival is more likely. Conversely, if the LV does not dilate and remains within the small to normal range, this indicates a poorer prognosis. In contrast the LVEF is not as reliable a predictor of outcome likely because SV is affected by these changes in EDV keeping the ratio (LVEF) constant or less predictive of outcome.

### Septic heart walls lose mass, increase percent water content and dilate

After bacterial challenge, we found through cMRI a significant progressive loss of LV total wall mass in survivors and non-survivors (Figure 3A). The LV steadily loses more weight in grams each day. The LV wall mass is ∼35 grams at baseline and survivors lose on average 1.7% +/- 0.6% of their LV wall mass/day. This loss is significantly more profound in non-survivors, with an average loss of 7.0% +/-1.7% per day. By 92 h, survivors have lost on average ∼2.5 grams (7.0%) of their LV wall mass. Accounting for the simple physical property of conservation of mass, a loss of total weight and a concurrent significant similar resultant percent increase in water content in survivors and non-survivors as measured by cMRI T2 (Figure 3B), means there is a decrease in dry mass in the LV wall.

**Figure 3.**
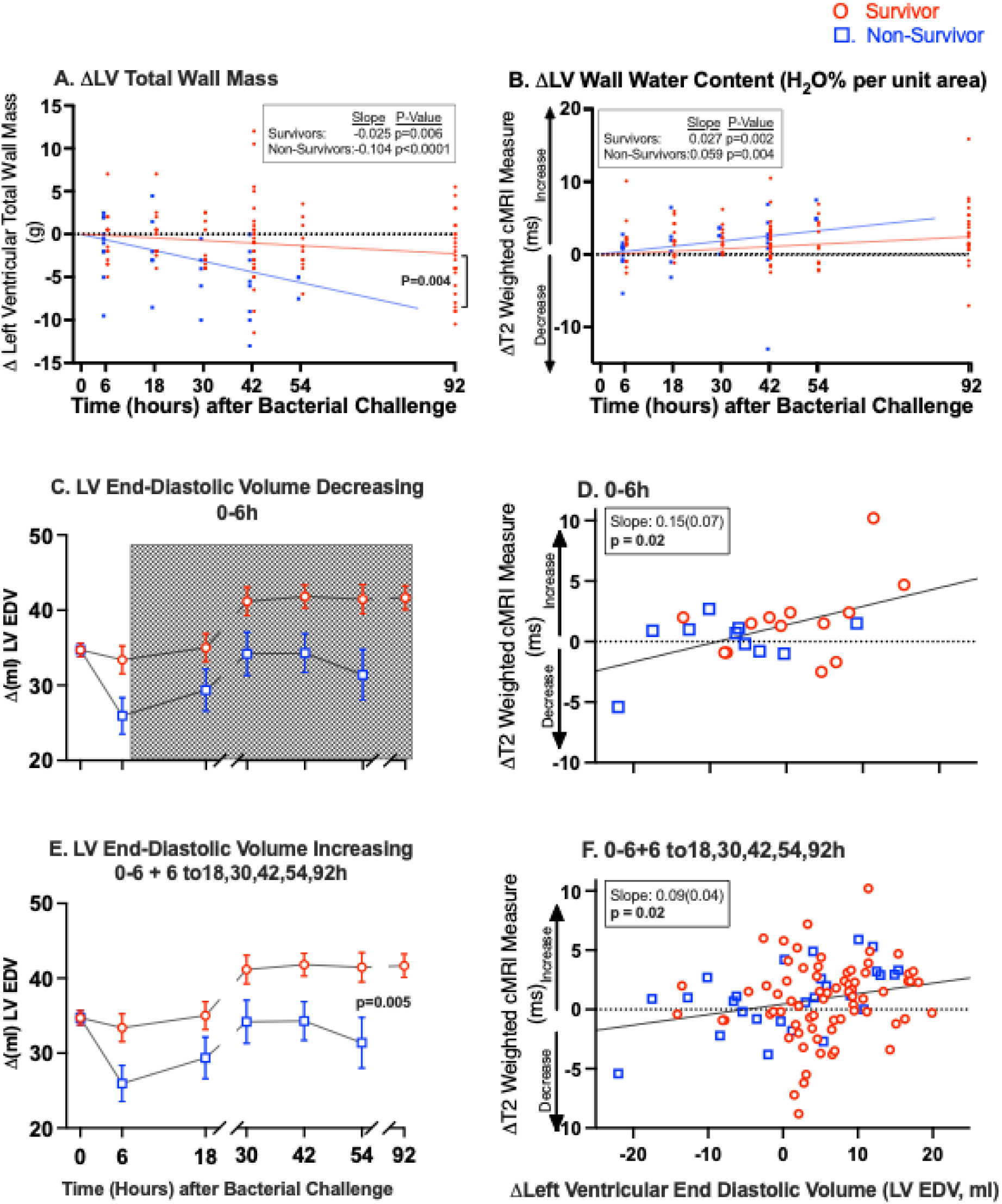
Septic Hearts Changes in LV Wall Mass, Water Content and EDV. LV total mass and water content parameters for surviving septic animals (red closed circles, red line) and non-surviving animal (blue closed circles, blue line) are plotted. LV wall mass Panel A and LV water content pre unit area of LV wall in Panel B. Using linear mixed models, a regression line was calculated for these individual values over time. A significant slope indicates an increase or decrease in each parameter for survivors or non-survivors and a comparison of the slopes was done to see if the two are different. Falls in LVEDV from 0 to 6 h after bacterial challenge are shown in Panel C. Association between cardiac edema and LVEDV from 0 to 6 h is shown Panel D. The changes in individual survivors’ values for EDV (open red circles) and non-survivors (open blue squares) are plotted on the x-axis verses the change in water per unit area of the LV wall on the y-axis. A significant positive slope indicates in survivors and non-survivors, the level of increasing edema in the left ventricular wall significantly correlates with the increase in end diastolic volume. Increasing changes in LV EDV from baseline to 92 h are shown in Panel E. In Panel F, similar to Panel D, shows the association between cardiac edema and LV EDV from baseline to 92h.

Progressive loss of LV total wall and dry mass over time likely leads to some wall thinning and ventricular chamber dilatation. Over time as total and dry myocardial mass declines, there is consequently a resultant significant percentage increase of water content within the LV wall, this effect potentially further enhances the wall thinning effect on LV EDV by increasing LV wall compliance. However, it is the resultant percent increase in water content (cMRI T2 increases) in the LV Wall (Figure 3C-F), and not the amount of mass loss (Figure 4A&B), that is closely and significantly associated with increases in LV EDV. Thus, indicating during sepsis how much of a resultant relative increase in the percent water content occurs in the LV wall with mass loss — and not how much mass is lost per se — is the critical factor associated with and pivotal to the increases in LV wall compliance and EDV. The resultant increases in percent water content and compliance in the ventricular wall seem to drive the stepwise ventricular dilation observed in both survivors and non-survivors from 0 to 30h. To the best of our knowledge, this mechanism of LV dilatation of on average 20-25% in our sepsis model [i.e., a decrease in total and dry mass with a resultant percentage increase in the LV wall water content (cMRI T2 increases), improving LV compliance] has not been described before.

**Figure 4.**
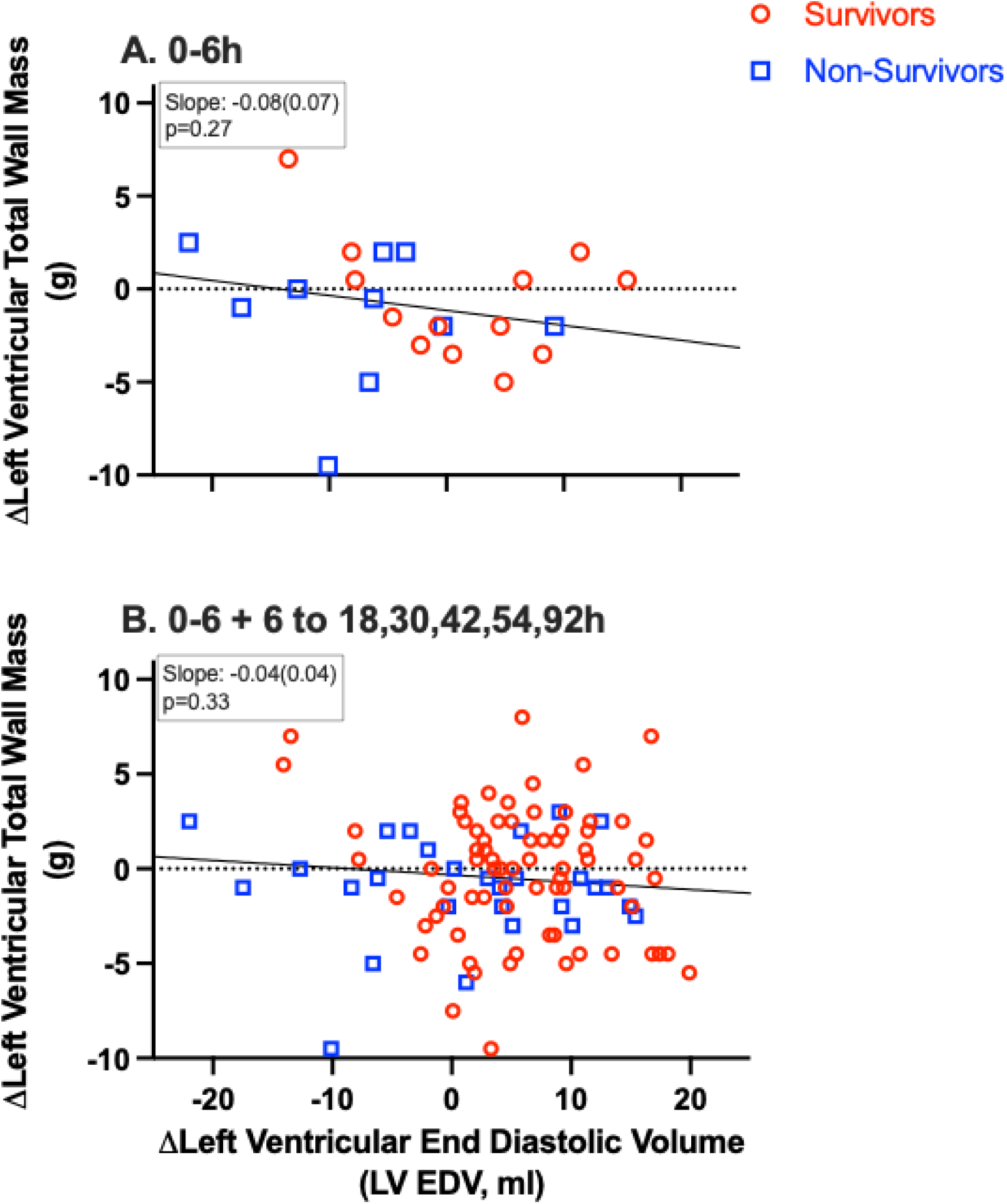
Septic Heart Changes in total mass and LVEDV. The changes in individual survivors’ values for EDV (open red circles) and non-survivors (open blue squares) are plotted on the x-axis verses the change total mass of the LV wall on the y-axis. In Panel A from 0-6h there was not a significant slope which indicates in survivors and non-survivors, the level of decreasing total wall mass in the left ventricular wall does not significantly correlate with the increase in end diastolic volume. Increasing changes in LV EDV from baseline to 92 h are shown in Panel B. Panel B, similar to Panel A, shows no association between cardiac total wall mass decrease and LV EDV from baseline to 92h.

By contrast, it has been more commonly found that, when LV wall percentage water content increases (cMRI T2 increases), total wall or dry mass remains constant or rises (rather than decreases, as it does in sepsis). This increase in edema (cMRI T2 increases) is a common finding in and around myocardial infarction (MIs) areas in the heart’s walls. In the case of MIs, these areas of increased edema in and around ischemic tissue decrease wall compliance (rather than increase compliance, as occurs in sepsis), reflecting a stiffer and less distensible myocardium.^11,15^

### Additional Supportive data from 0-6h, bolstering the hypothesis that the resultant percent increases in relative water content in the LV wall during mass loss causes the increases in LV compliance and EDV

#### Baseline to Six hours, the LV EDV decreases in septic animals (stage one injury)

Further supporting our “compliance” hypothesis during sepsis, were the relationships we found between total mass loss, cMRI T2, and LV EDV from time 0-6h post bacterial challenge. This is when the LV compliance injury is occurring, and the LV EDV is overall declining (Figure 3C). Beginning from time 0-6h, significant cardiac total mass loss was not significantly associated with increases in LV EDV (Figure 4A). However, this mass loss was accompanied by significant increases in cMRI T2, which did in turn significantly correlate with increases in LV EDV (Figure 3D). This implies cMRI T2 serves as an index of myocardial compliance: during mass loss, the greater the resultant increase in T2, (i.e., larger the concomitant increase in LV wall water content), the greater the increase in EDV. Importantly, even though overall EDV declines between 0-6h during this injurious period in the LV wall, the myocardium’s with the most pronounced water-percent increases from baseline (i.e., have a greater and greater percent increase in water content as mass is lost 0-6h) exhibit the greatest increases in compliance and, consequently, the largest relative rises in EDV – demonstrating that the water-content-compliance relationship holds true even as EDV falls during this injurious interval to the LV wall.

#### Combined 0 to 6h injury stage one, and stage two 6 to 18,30,54 and 92h when compliance of the LV wall overall is increasing

From 0 to 92h after bacterial challenge, we observed early decreases in EDV and later progressive increases in LV EDV. However, throughout the study, LV wall water percentage content progressively increased from 0 to 92h alongside a progressive significant decline in total myocardial mass, leading to a significant rise in the percent water-to-dry mass per unit area of myocardial tissue in both survivors (Figure 3E) and non-survivors alike (i.e., cMRI T2 measurements significantly increases). During this entire time from 0 to 92h the cMRI T2 increases (i.e., percent water increases in the LV wall) in the LV wall were associated with increases in EDV. In contrast the LV total mass loss did not significantly correlate with changes in EDV throughout the study (Figure 4A&B). Thus, the resultant percent increase in water content with dry mass loss is likely the critical factor that enhances wall compliance and drives ventricular dilation, thereby elevating LV EDV (Figure 3E&F). In a previous study, non-septic controls that were treated identically, except for receiving intrabronchial saline instead of *S.aureus*, demonstrated no significant changes in LV total wall mass or percent water content in the LV over 92h.^7^

### Myocyte and mitochondria ultrastructural damage is associated with mortality, with earlier deaths exhibiting greater capillary ultrastructure processing of damaged components

#### Light Microscopy

Light microscopy revealed only minimal inflammatory cell infiltrate and interstitial edema in both survivors and non-survivors as found in our previous studies.^7,16^

#### Capillary Injury and ultrastructure processing of damaged components in the LV wall

LV tissue obtained from the seven early non-survivors (9-28h after bacterial challenge) showed the presence of vesiculo-vacuolar organelles and pericapillary particle deposits which were consistently graded as mild to moderate (Figure 5A, see Table S1 for individual animal’s time-of-death and scores). This is evidence of ongoing ultrastructure processing of the damaged components within the LV wall in the interstitial space around the microvasculature in early deaths. This was significantly increased when compared to the five non-survivors that died late (from 44-83h), suggesting there was more damage in some LV tissues at this early time point and perhaps more attempts at early repair (Figure 5A). Thereafter, this summary score of both late-stage septic non-survivors from 44-83h and septic survivors sacrificed at 66h (a comparable time point) did not differ significantly, indicating no further associated differences associated with survival in capillary processing of damaged components in the LV wall at this later timepoint (Figure 5A). The mean endothelial cell score of mild to moderate increased swelling was not significantly different comparing survivors or early deaths to late deaths, suggesting that ultrastructure injury of capillaries is not associated closely with outcome (Figure 5A).

**Figure 5.**
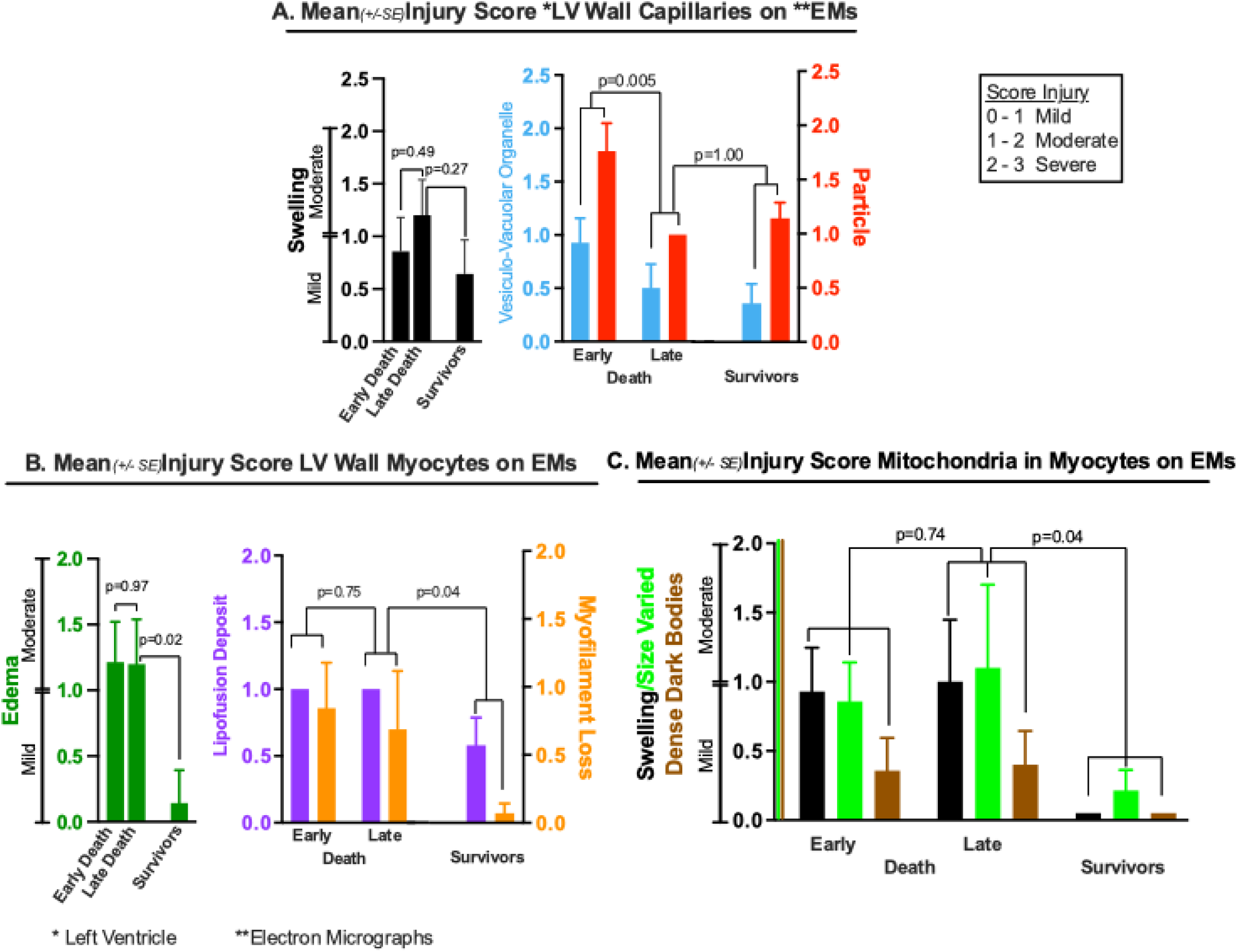
Septic Heart LV wall Ultrastructure Changes. Graphical illustration of Table S1. Panel A compares the mean injury score calculated by EM in the capillaries of the LV walls between early and late deaths, and survivors. The scores in Panel A shows capillary cell swelling (right graph) vesiculo-vacuolar organelles in the endothelial cells, (blue graph) and particles (red graph). Panel B compares the mean injury scores calculated by EM within the myocytes between early and late deaths, and survivors. The scores in Panel B shows myocyte intracellular edema (right graph) lipofusion deposits within the myocytes, (purple graph) and myofilament loss (orange graph). Panel C compares the mean injury scores/morphological changes calculated by EM within the mitochondria of the myocytes between early and late deaths, and survivors. The scores in Panel C shows mitochondrial swelling (closed graph) variation of sizes/fusion, (green graph) and black dot deposits (brown graph).

Myocyte damage mean scores on Ems — as measured by edema, lipofuscin deposits, myofilament fragmentation — and myocyte mitochondrial damage (as measured by mitochondrial swelling, variation in sizes, and the presence of dark dense bodies) was similarly moderately significantly increased in the 12 of the 18 non-survivors that died early and late for which we were able to obtain LV tissues (Figure 5B&C). However, in contrast to capillaries, these damages to myocytes and their mitochondria mean scores were significantly worse in the five non-survivors that died late between 44-83h compared to the 7 survivors euthanized at a comparable timepoint 66h (Figure 5B&C and Table S1). The typical finding in septic animals at the time of death on EMs within around endothelial cells of the microvasculature (i.e., the capillaries) are shown in Figure 6A-F and for myocytes and their mitochondria the typical ultrastructural changes are shown in Figure 6G-N.

**Figure 6.**
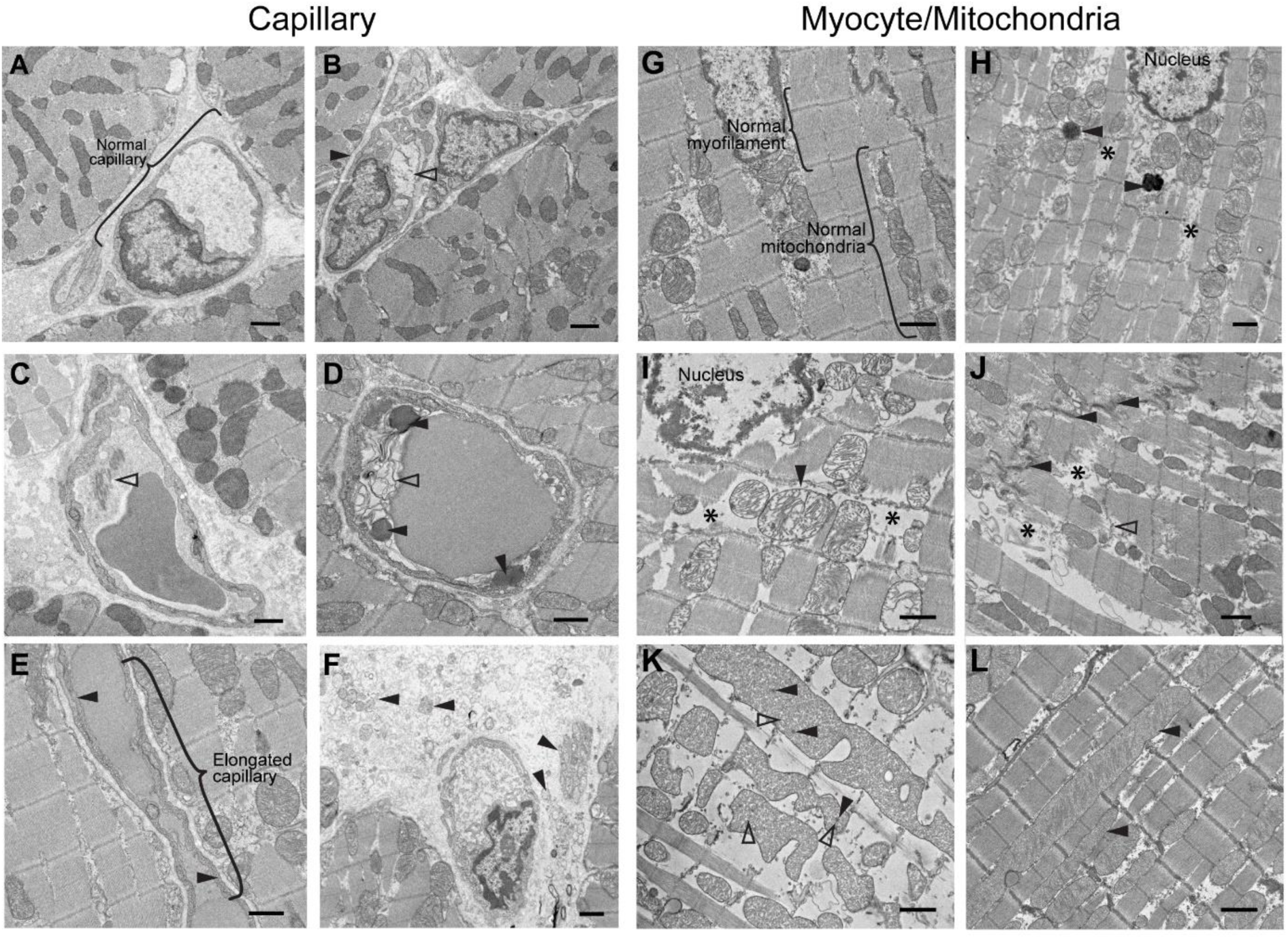
Septic Heart LV wall Ultrastructure Changes. Representative Electron Micrographs. In Panel A, a micrograph a normal capillary in the left ventricular wall. Panel B, a partially damaged capillary endothelial cell. Note that the endothelial cell on the right side clearly shows edema (open arrowheads) whereas the endothelial cell on the left side shows normal density (closed arrowheads). Panel C, a capillary, contains fibrin like material (open arrowhead). Panel D, multiple dense bodies (closed arrowheads) and organelles (open arrowhead) in the capillary lumen. Panel E, an elongated capillary endothelial cell shows vesiculo-vacuolar organelle in the endothelial cell wall (closed arrowheads). Panel F, fragments of tissues in the interstitial space near capillary (closed arrowheads). Panel G, an intact myocyte with normal ultrastructure. Panel H, myocyte with mild intracellular edema and myofilament loss (*), and lipofuscin (closed arrowheads). Panel I, myocyte with severe edema, myofilament loss (*) and mitochondria swelling (closed arrowhead). Panel J, severe myofilament loss and edema (*) near disrupted intercalated discs (closed arrowheads) and z-line disruption of myocytes (open arrowhead). Panel K, mitochondria with varied morphological changes, included various fusion sizes (open arrowheads) and black dot deposits (closed arrowheads). Panel L, giant mitochondria were observed (closed arrowheads). Bars equal one micron for all panels.

Previously we compared septic large animals with E. coli peritonitis sacrificed at 48h LV tissue to non-septic controls sacrificed at 48h.^16^ Our findings here, of significantly increased diffuse endothelial cell edema and non-occlusive fibrin deposition on EMs as well as myofibril disruption and loss, were also found to be significantly increased on EMs in septic animals at 48h when compared to non-septic controls.^16^ We demonstrate here a sepsis-induced greater ultrastructure injury in non-survivors compared to survivors in cardiac myocytes and their mitochondria. The endothelial cells appear to be markedly involved in processing damaged tissue components from myocytes. By contrast, the damage present in capillaries is not as significantly associated with outcome as the damage in myocytes. The more severe damage to myocytes in non-survivors than survivors is dispersed through the LV wall, potentially impacting myofilaments three different ways and thus causing the drop in EF and more severe decrease in EDV: 1) edema in the myocyte is hindering contraction and relaxation; 2). myofilament fragmentation is weakening the contractile and relaxation apparatus; and 3) mitochondrial disruption is making processing energy sources inefficient, retarding contraction and relaxation. These injuries to the myocytes and their mitochondria and increased processing of damaged LV components in capillaries, are shown here — for the first time, to the best of our knowledge — to be part of the septic cardiomyopathy and more extensive in non-survivors at early and late time points of death than in survivors (Figure 4A-C and Table S1). These occur without evidence of in survivors or non-survivors of prominent cellular inflammatory infiltrates, tissue ischemia, bacterial remnants, or signs of developing fibrosis on light microscopy.

### Multi-Organ Injury Mirrors Cardiac Outcomes

The consequences of septic shock clinically are not isolated to the heart but commonly lead to multi-organ failure that is the prelude to mortality.^17^ We observed this result in our animal model with significant injury in non-survivors compared to survivors across several organ systems: i) kidney function as measured by mean serum creatinine over the entire study period (Figure S2A-C); ii) liver function as measured by mean aspartate aminotransferase and alanine transaminase elevations from 48 to 72 h (Figure S2D&E); iii) metabolic function as measured by mean pH at 18 to 24h (Figure S3A); iv) pulmonary function as measured by the ratio of arterial oxygen content over fractional inspired oxygen concentration from 18 to 42h (Figure S3B).

Other parameters’ mean values showed no meaningful differences between survivors and non-survivors over the entire study: i) other blood gas measurements (Figure S3); ii) myocardial perfusion as measured by cMRI (Figure S4); iii) serum cytokines (Figure S5-S6); iv) complete blood cell counts (Figure S7); and v) serum chemistries (Figure S8-S9).

## Discussion

### The early injury stage at 0 to 6h

Using our large-animal septic shock model that replicates human septic cardiomyopathy,^2,4,5^ we investigated whether temporal successive alterations in LV wall ultrastructure’s, water content, and mass could explain the basis of serial LVEF and EDV changes associated with outcomes. Maximal diastolic dysfunction occurred within 6h of bacterial challenge, marked by a steep decline in LV compliance and LVEDV – significantly greater in non-survivors, and sustained for the full 5-day study. Due to the time needed to develop and present clinically with septic shock in humans, little may be reported about what happens very early in the development. Despite this, non-survivors have been found to demonstrate a pattern of diastolic filling suggestive of abnormal left ventricular relaxation compared with survivors within 24 hours of septic shock.^18–20^ Furthermore, in our animal model, EMs at time of death showed more severe myocyte damage in non-survivors compared to survivors. This included increased edema of myocytes, myofilament fragmentation, lipofuscin deposits, and abnormal myocyte mitochondria swelling.

The finding of ultrastructure damage appears to be consistent across different species and sepsis models. Vanasco et al. showed that 6h after LPS injection in a murine model, there were significant increases in mitochondrial number, size, and damage, a finding echoed in a LPS model of felines.^21,22^ Celes et al. further showed disruption to the contractile apparatus, especially loss of dystrophin, at 6h in a peritoneal model of murine septic shock.^23^ Such ultrastructural damage can explain why there is compromise in the myofilaments’ ability to “relax”, restricting LV dilation and contraction and sustaining a persistently reduced EDV in non-survivors until death. Additionally, we found on EM more vesiculo-vaculor organelles and particles in and around capillaries which were associated with earlier deaths, suggesting the occurrence of increased processing of severely damaged LV wall ultrastructure.

### Passive recovery of diastolic function (6-30h)

After 6h, LV compliance increases, leading to a steady rise in LV EDVs in both survivors and non-survivors. The correlation between higher LVEDV and survival corroborates with our and others previous findings in septic patients and animal models.^1,2,20,22,24^ During this early recovery phase, cMRI T2 revealed parallel increases in percent water content in the LV walls in survivors and non-survivors. Notably, non-survivors exhibited significantly greater mass loss, likely reflecting more extensive tissue injury. However, this mass loss did not correlate with changes in EDV (Figure 4AB). Instead, the resultant increase in percent relative water content with mass loss was significantly associated with increases in EDV and appeared to serve as a surrogate marker for improving LV compliance (Figure 3C-F). While increased myocardial edema is generally suggestive of poorer prognosis in myocarditis and myocardial infarctions, in other scenarios such as sudden cardiac arrest — and now sepsis — the presence of myocardial edema was found to be favorable, reflecting return of diastolic function.^25,26^ In different pathophysiological mechanisms of disease, edema or increase in tissue water content has different implications. A one-size solution/theory does not fit all.

Our findings indicate that loss of dry mass raises relative percent water content, enhancing LV wall compliance, and facilitates the LV dilation in both survivors and non survivors. However, we observed significantly greater mass loss in non-survivors, and on EMs more myocyte edema and myofilament fragmentation, corresponding to lower absolute EDV throughout the study. By contrast, survivors suffered significantly less structural damage on EMs and less decreases in EDV, and as mass was lost, the relative increase in LV water content resulted in development of a “super” compliance causing a more robust absolute recovery in EDV. Prior murine studies show evolving myocardial cellular changes during sepsis, including altered contractility, transiently elevated ATPase activity, disrupted calcium handling, and mitochondrial dysfunction.^27,28^ Similarly, Takasu et al. reported EM findings in 17 septic human non-survivors (medium 6 days from diagnosis till death) showing myocyte injury consistent with our observations as well as no evidence of cell death by apoptosis.^29^ Together, this suggests that passive functional recovery is underpinned by active subcellular repair processes removing damaged components of cells increasing relative water content in LV wall, compliance and EDV. In survivors, there is less ultrastructure damage and/or better repair and with increased relative LV wall water content developing “super” compliance to the LV wall and LV dilation to 20 to 25% greater than baseline. Non-survivors have greater persistent ultrastructure injury, worse repair, never-ending decreased compliance and inability to increase EDV from 6 to 30h greater than baseline.

### Active recovery of systolic function (Beyond 30h)

An active repair process occurs within the LV wall, marked by ongoing mass loss likely representing clearance of damaged cellular components, beginning soon after septic injury and persisting for over 5 days. EMs revealed that there are organelles, dense bodies, vesicles, and vacuoles within and surrounding microcirculatory endothelial cells, indicating ongoing clearance of damaged subcellular structures. In septic human non-survivors, late in their clinical course, EMs showed diffusely scattered foci of actin and myosin disruption, consistent with ongoing myofilament lysis despite attempts at repair.^30^ Systolic function (LVEF) in humans and our animal models takes 10–14 days post-septic shock onset^1,2^ for full recovery (LVEF ∼55–65%). Mass loss with functional recovery has been observed in human CMR studies of myocarditis, where myocardial mass decreased at a 6-month follow-up as function improved.^31^ We propose that prolonged systolic recovery reflects the need for active structural remodeling, including repair of myofilament fragmentations and resolution of myocyte edema and mitochondrial swelling. These restorative processes contribute to recovery of contractile apparatus involved in systolic function. Progressively, myofilaments contraction improves over the ensuing days. With complete return to normal pre sepsis levels of contraction function in survivors by 10 to 14 days.^1,2^

### Systemic Inflammation is the cause of reversible Myocyte ultrastructure injury

We previously demonstrated that a variety of inflammatory insults can all reproduce the same cardiac dysfunction pattern in large animal models. This included the intravenous administration of human tumor necrosis factor and bacterial endotoxin,^32^ as well as intraperitoneal and bronchial challenges of viable and non-viable gram-positive and gram-negative bacteria.^5,33^ We found that, across different bacterial doses,^24^ bacterial species, bacterial viability^33^, type of inflammatory mediator and routes^34^ of administration, each challenge consistently reproduced the same pattern of cardiac injury characteristic of the septic cardiomyopathy in humans. This suggests an array of structurally and functionally different nonspecific systemic inflammatory triggers can induce this same phenotype of cardiomyopathy.

This finding supports the hypothesis that septic cardiomyopathy arises from a source in the body that can activate a nonspecific systemic diffuse inflammation-mediated immune response, regardless of the nature or site of the source of inflammation trigger. Consistent with this notion, cytokine exposure to isolated ventricular cardiomyocytes of guinea pigs was found to exert a negative inotropic effect.^35^ Further, inhibition of inflammatory pathways with a nonselective tyrosine kinase inhibitor attenuated the decline in EF in septic cardiomyopathy, reducing multiorgan injury, and significantly improved survival in our animal model of sepsis.^36^ These results highlight the central role of inflammation in the initial pathogenesis of septic cardiomyopathy.

### Exclusion of Likely Alternative Mechanisms

Two frequently proposed alternative mechanisms for septic cardiomyopathy are: 1) diffuse ischemic tissue injury and 2) high-dose catecholamine stress-induced injury.^37–41^ Serial stress perfusion cMRIs done here demonstrated preserved or even enhanced microcirculatory blood flow in both survivors and non-survivors throughout the 5-day study (Figure S4). This observation is inconsistent with myocardial tissue ischemia as the primary etiology of the septic cardiomyopathy. Second, there is no evidence in our model — on light microscopy or EM imaging — of ischemic injury to myocardial cells or occlusive injury to the microcirculation. Moreover, serial plasma troponin I levels, a sensitive marker of myocyte ischemia and death, were found not to be elevated in previous studies.^6^ In both humans and in our septic model, coronary sinus blood flow during this injury is not impaired.^16,42,43^ Further, in models where catecholamine release was suppressed and no exogenous catecholamines were administered, the typical cardiac injury still occurred.^6^ These results argue against ischemic or catecholaminergic mechanisms as primary drivers of septic cardiomyopathy.

## Conclusion

Septic cardiomyopathy is an acute, diffuse, non-ischemic, non-stress-induced inflammatory injury affecting the ultrastructure of cardiac microcirculatory endothelial cells and myocytes. This injury rapidly follows bacterial challenge with maximal loses in ventricular compliance, chamber filling, and EDV occurring very early on — within 6h — along with decreases in EF. Noteworthy, these findings imply that the critical cardiac injury in sepsis occurs early and may be largely complete before patients present clinically, potentially explaining the limited efficacy of late anti-inflammatory or non-antibiotic interventions in human sepsis trials.^44,45^ Myocardial repair begins early, involving removal of damaged subcellular components. As damaged ultrastructure are removed, the LV wall thins, the percent water content increases in the LV wall increasing compliance, and the ability of ventricles to dilate returns to normal or “supernormal” by 30h. This marks a passive mechanism of recovery of diastolic function. Later, as dry mass loss continues over days, myofilament contraction improves with the EF returning to normal in progressive stages. This suggests an active repair mechanism to edematous myocytes and disrupted myofilaments and swollen myocyte mitochondria over 14 days.^1,2^ Importantly, non-survivors experience greater severity of ultrastructural damage, insufficient repair, and persistently depressed LVEDV, representing critical distinctions between survival and non-survivable injury. The LVEDV, and not the LVEF, appears to be an excellent pathophysiological biomarker and indicator of the degree of ultrastructure damage and eventual outcomes from septic shock. Our observations extend beyond the heart: Non-survivors also exhibited more severe injury in other organs (kidneys, liver, lungs, metabolic function) over a similar course, consistent with a systemic inflammatory response leading to multi-organ failure from ultrastructure damage. Thus, the myocardial injury in septic shock should not be viewed in isolation, but as part of a broader syndrome of inflammation-induced multi-organ dysfunction and serial changes in EDV — a pathophysiological biomarker and indicator of the degree of ultrastructure damage affecting multi-organ failure and sepsis outcomes.

## Author Contributions

Designing research studies: CN, VJF, SBS, WNF, MAS, RLD, HGK

Conducting experiments CN, SBS, VJF, JF, MYF, WNA

Acquiring data CN, Z-XY, VJF, SBS, MYC, JW

Analyzing data CN, JF, Z-XY, YL, JW, MYC,VJF

Providing reagents PT-P

Writing the manuscript: CN, VJF, WNF, HGK, ZNS, Z-XY, IC-P, MAS, RLD

Writing – review & editing: CN, VJF, SBS, JS, WNF, JW, ICP, MAS, RLD, HGK

## Acknowledgements

**Funding:** This work was supported by NIH intramural funding from the NIH Clinical Center.

**Role of funding source:** The work by the authors was conducted as part of US government– funded research; however, the opinions expressed are not necessarily those of the National Institutes of Health (NIH).

## Supplementary Materials

Methods

Limitations

Figures S1 to S9

Table S1

References #6, 7, 16, 46, 47

## Abbreviations

AUC: area under the curve
cMRI: cardiac magnetic resonance images
CI: Cardiac Index
CVP: central venous pressure
ESV: end systolic function
LV: left ventricular
EDV: end diastolic volume
EF: ejection fraction
EM: electron micrographs
HR: heart rate
ICU: intensive care units
LVEF: left ventricular ejection fraction
NIH: National Institutes of Health
PAOP: pulmonary artery occlusion pressure
PVRI: pulmonary vascular resistance index
ROC: receiver operating characteristic
SV: stroke volume
SVRI: systemic vascular resistance index

## References

1. Parker MM, Shelhamer JH, Bacharach SL, Green MV, Natanson C, Frederick TM, Damske BA, Parrillo JE. Profound but reversible myocardial depression in patients with septic shock. Ann Intern Med. 1984;100:483–490. doi: 10.7326/0003-4819-100-4-483

2. Natanson C, Fink MP, Ballantyne HK, MacVittie TJ, Conklin JJ, Parrillo JE. Gram-negative bacteremia produces both severe systolic and diastolic cardiac dysfunction in a canine model that simulates human septic shock. J Clin Invest. 1986;78:259–270. doi: 10.1172/jci112559

3. Fan D, Wu R. Mechanisms of the septic heart: From inflammatory response to myocardial edema. J Mol Cell Cardiol. 2024;195:73–82. doi: 10.1016/j.yjmcc.2024.08.003

4. Minneci PC, Deans KJ, Hansen B, Parent C, Romines C, Gonzales DA, Ying SX, Munson P, Suffredini AF, Feng J, et al. A canine model of septic shock: balancing animal welfare and scientific relevance. Am J Physiol Heart Circ Physiol. 2007;293:H2487–2500. doi: 10.1152/ajpheart.00589.2007

5. Solomon SB, Wang D, Sun J, Kanias T, Feng J, Helms CC, Solomon MA, Alimchandani M, Quezado M, Gladwin MT, et al. Mortality increases after massive exchange transfusion with older stored blood in canines with experimental pneumonia. Blood. 2013;121:1663–1672. doi: 10.1182/blood-2012-10-462945

6. Ford VJ, Applefeld WN, Wang J, Sun J, Solomon SB, Klein HG, Feng J, Lertora J, Parizi-Torabi P, Danner RL, et al. In a Canine Model of Septic Shock, Cardiomyopathy Occurs Independent of Catecholamine Surges and Cardiac Microvascular Ischemia. J Am Heart Assoc. 2024;13:e034027. doi: 10.1161/jaha.123.034027

7. Ford VJ, Applefeld WN, Wang J, Sun J, Solomon SB, Sidenko S, Feng J, Sheffield C, Klein HG, Yu ZX, et al. Cardiac Magnetic Resonance Studies in a Large Animal Model That Simulates the Cardiac Abnormalities of Human Septic Shock. J Am Heart Assoc. 2024;13:e034026. doi: 10.1161/jaha.123.034026

8. Schmittinger CA, Dünser MW, Torgersen C, Luckner G, Lorenz I, Schmid S, Joannidis M, Moser P, Hasibeder WR, Halabi M, et al. Histologic Pathologies of the Myocardium in Septic Shock: A Prospective Observational Study. Shock. 2013;39:329–335. doi: 10.1097/SHK.0b013e318289376b

9. Vasques-Nóvoa F, Laundos TL, Madureira A, Bettencourt N, Nunes JPL, Carneiro F, Paiva JA, Pinto-do-Ó P, Nascimento DS, Leite-Moreira AF, et al. Myocardial Edema: an Overlooked Mechanism of Septic Cardiomyopathy? Shock. 2020;53:616–619. doi: 10.1097/shk.0000000000001395

10. Augusto JB, Nordin S, Vijapurapu R, Baig S, Bulluck H, Castelletti S, Alfarih M, Knott K, Captur G, Kotecha T, et al. Myocardial Edema, Myocyte Injury, and Disease Severity in Fabry Disease. Circ Cardiovasc Imaging. 2020;13:e010171. doi: 10.1161/circimaging.119.010171

11. Bonfig NL, Soukup CR, Shah AA, Olet S, Davidson SJ, Schmidt CW, Peterson R, Henry TD, Traverse JH. Increasing myocardial edema is associated with greater microvascular obstruction in ST-segment elevation myocardial infarction. Am J Physiol Heart Circ Physiol. 2022;323:H818–h824. doi: 10.1152/ajpheart.00347.2022

12. Havaldar AA. Evaluation of sepsis induced cardiac dysfunction as a predictor of mortality. Cardiovascular Ultrasound. 2018;16:31. doi: 10.1186/s12947-018-0149-4

13. Lee KJ, Kim YK, Jeon K, Ko RE, Suh GY, Oh DK, Lim SY, Lee YJ, Lee SY, Park MH, et al. Shock indices are associated with in-hospital mortality among patients with septic shock and normal left ventricular ejection fraction. PLoS One. 2024;19:e0298617. doi: 10.1371/journal.pone.0298617

14. Yamamoto K, Masuyama T, Tanouchi J, Doi Y, Kondo H, Hori M, Kitabatake A, Kamada T. Effects of heart rate on left ventricular filling dynamics: assessment from simultaneous recordings of pulsed Doppler transmitral flow velocity pattern and haemodynamic variables. Cardiovasc Res. 1993;27:935–941. doi: 10.1093/cvr/27.6.935

15. Cross CE, Rieben PA, Salisbury PF. Influence of coronary perfusion and myocardial edema on pressure-volume diagram of left ventricle. Am J Physiol. 1961;201:102–108. doi: 10.1152/ajplegacy.1961.201.1.102

16. Solomon MA, Correa R, Alexander HR, Koev LA, Cobb JP, Kim DK, Roberts WC, Quezado ZM, Scholz TD, Cunnion RE, et al. Myocardial energy metabolism and morphology in a canine model of sepsis. Am J Physiol. 1994;266:H757–768. doi: 10.1152/ajpheart.1994.266.2.H757

17. Sun GD, Zhang Y, Mo SS, Zhao MY. Multiple Organ Dysfunction Syndrome Caused by Sepsis: Risk Factor Analysis. Int J Gen Med. 2021;14:7159–7164. doi: 10.2147/ijgm.S328419

18. Munt B, Jue J, Gin K, Fenwick J, Tweeddale M. Diastolic filling in human severe sepsis: An echocardiographic study. Critical Care Medicine. 1998;26:1829–1833.

19. Ehrman RR, Bredell BX, Harrison NE, Favot MJ, Haber BD, Welch RD, Levy PD, Sherwin RL. Increasing illness severity is associated with global myocardial dysfunction in the first 24 hours of sepsis admission. Ultrasound J. 2022;14:32. doi: 10.1186/s13089-022-00282-6

20. Landesberg G, Gilon D, Meroz Y, Georgieva M, Levin PD, Goodman S, Avidan A, Beeri R, Weissman C, Jaffe AS, et al. Diastolic dysfunction and mortality in severe sepsis and septic shock. European Heart Journal. 2011;33:895–903. doi: 10.1093/eurheartj/ehr351

21. Vanasco V, Saez T, Magnani ND, Pereyra L, Marchini T, Corach A, Vaccaro MI, Corach D, Evelson P, Alvarez S. Cardiac mitochondrial biogenesis in endotoxemia is not accompanied by mitochondrial function recovery. Free Radic Biol Med. 2014;77:1–9. doi: 10.1016/j.freeradbiomed.2014.08.009

22. Joshi MS, Julian MW, Huff JE, Bauer JA, Xia Y, Crouser ED. Calcineurin regulates myocardial function during acute endotoxemia. Am J Respir Crit Care Med. 2006;173:999–1007. doi: 10.1164/rccm.200411-1507OC

23. Celes MRN, Torres-Dueñas D, Malvestio LM, Blefari V, Campos EC, Ramos SG, Prado CM, Cunha FQ, Rossi MA. Disruption of sarcolemmal dystrophin and β-dystroglycan may be a potential mechanism for myocardial dysfunction in severe sepsis. Laboratory Investigation. 2010;90:531–542. doi: 10.1038/labinvest.2010.3

24. Natanson C, Danner RL, Fink MP, MacVittie TJ, Walker RI, Conklin JJ, Parrillo JE. Cardiovascular performance with E. coli challenges in a canine model of human sepsis. Am J Physiol. 1988;254:H558–569. doi: 10.1152/ajpheart.1988.254.3.H558

25. Zorzi A, Susana A, De Lazzari M, Migliore F, Vescovo G, Scarpa D, Baritussio A, Tarantini G, Cacciavillani L, Giorgi B, et al. Diagnostic value and prognostic implications of early cardiac magnetic resonance in survivors of out-of-hospital cardiac arrest. Heart Rhythm. 2018;15:1031–1041. doi: 10.1016/j.hrthm.2018.02.033

26. Zorzi A, Mattesi G, Baldi E, Toniolo M, Guerra F, Cauti FM, Cipriani A, De Lazzari M, Muser D, Stronati G, et al. Prognostic Role of Myocardial Edema as Evidenced by Early Cardiac Magnetic Resonance in Survivors of Out-of-Hospital Cardiac Arrest: A Multicenter Study. Journal of the American Heart Association. 2021;10:e021861. doi: 10.1161/JAHA.121.021861

27. Powers FM, Farias S, Minami H, Martin AF, Solaro RJ, Law WR. Cardiac myofilament protein function is altered during sepsis. J Mol Cell Cardiol. 1998;30:967–978. doi: 10.1006/jmcc.1998.0661

28. Joseph LC, Kokkinaki D, Valenti MC, Kim GJ, Barca E, Tomar D, Hoffman NE, Subramanyam P, Colecraft HM, Hirano M, et al. Inhibition of NADPH oxidase 2 (NOX2) prevents sepsis-induced cardiomyopathy by improving calcium handling and mitochondrial function. JCI Insight. 2017;2. doi: 10.1172/jci.insight.94248

29. Takasu O, Gaut JP, Watanabe E, To K, Fagley RE, Sato B, Jarman S, Efimov IR, Janks DL, Srivastava A, et al. Mechanisms of cardiac and renal dysfunction in patients dying of sepsis. Am J Respir Crit Care Med. 2013;187:509–517. doi: 10.1164/rccm.201211-1983OC

30. Rossi MA, Celes MRN, Prado CM, Saggioro FP. MYOCARDIAL STRUCTURAL CHANGES IN LONG-TERM HUMAN SEVERE SEPSIS/SEPTIC SHOCK MAY BE RESPONSIBLE FOR CARDIAC DYSFUNCTION. Shock. 2007;27:10–18. doi: 10.1097/01.shk.0000235141.05528.47

31. Aquaro Giovanni D, Ghebru Habtemicael Y, Camastra G, Monti L, Dellegrottaglie S, Moro C, Lanzillo C, Scatteia A, Di Roma M, Pontone G, et al. Prognostic Value of Repeating Cardiac Magnetic Resonance in Patients With Acute Myocarditis. JACC. 2019;74:2439–2448. doi: 10.1016/j.jacc.2019.08.1061

32. Natanson C, Eichenholz PW, Danner RL, Eichacker PQ, Hoffman WD, Kuo GC, Banks SM, MacVittie TJ, Parrillo JE. Endotoxin and tumor necrosis factor challenges in dogs simulate the cardiovascular profile of human septic shock. J Exp Med. 1989;169:823–832. doi: 10.1084/jem.169.3.823

33. Natanson C, Danner RL, Elin RJ, Hosseini JM, Peart KW, Banks SM, MacVittie TJ, Walker RI, Parrillo JE. Role of endotoxemia in cardiovascular dysfunction and mortality. Escherichia coli and Staphylococcus aureus challenges in a canine model of human septic shock. J Clin Invest. 1989;83:243–251. doi: 10.1172/jci113866

34. Eichenholz PW, Eichacker PQ, Hoffman WD, Banks SM, Parrillo JE, Danner RL, Natanson C. Tumor necrosis factor challenges in canines: patterns of cardiovascular dysfunction. Am J Physiol. 1992;263:H668–675. doi: 10.1152/ajpheart.1992.263.3.H668

35. Sugishita K, Kinugawa K-i, Shimizu T, Harada K, Matsui H, Takahashi T, Serizawa T, Kohmoto O. Cellular Basis for the Acute Inhibitory Effects of IL-6 and TNF-α on Excitation-contraction Coupling. Journal of Molecular and Cellular Cardiology. 1999;31:1457–1467. doi: 10.1006/jmcc.1999.0989

36. Sevransky JE, Shaked G, Novogrodsky A, Levitzki A, Gazit A, Hoffman A, Elin RJ, Quezado ZMN, Freeman BD, Eichacker PQ, et al. Tyrphostin AG 556 improves survival and reduces multiorgan failure in canine Escherichia coli peritonitis. The Journal of clinical investigation. 1997;99 8:1966–1973.

37. Cappelletti S, Ciallella C, Aromatario M, Ashrafian H, Harding S, Athanasiou T. Takotsubo Cardiomyopathy and Sepsis. Angiology. 2017;68:288-303. doi: 10.1177/0003319716653886

38. Jing C, Wang Y, Kang C, Dong D, Zong Y. Clinical features of patients with septic shock-triggered Takotsubo syndrome: a single-center 7 case series. BMC Cardiovascular Disorders. 2022;22:340. doi: 10.1186/s12872-022-02787-3

39. Hahn PY, Wang P, Tait SM, Ba ZF, Reich SS, Chaudry IH. Sustained elevation in circulating catecholamine levels during polymicrobial sepsis. Shock. 1995;4:269–273. doi: 10.1097/00024382-199510000-00007

40. Ammann P, Fehr T, Minder E, Günter C, Bertel O. Elevation of troponin I in sepsis and septic shock. Intensive Care Medicine. 2001;27:965–969. doi: 10.1007/s001340100920

41. Mehta NJ, Khan IA, Gupta V, Jani K, Gowda RM, Smith PR. Cardiac troponin I predicts myocardial dysfunction and adverse outcome in septic shock. International Journal of Cardiology. 2004;95:13–17. doi: 10.1016/j.ijcard.2003.02.005

42. Dhainaut J-F, Marin N, Chiche J. Coronary circulation in sepsis. Coronary Circulation and Myocardial Ischemia. 2000:36–45.

43. Cunnion RE, Schaer GL, Parker MM, Natanson C, Parrillo JE. The coronary circulation in human septic shock. Circulation. 1986;73:637–644. doi: 10.1161/01.cir.73.4.637

44. Sweeney DA, Danner RL, Eichacker PQ, Natanson C. Once is not enough: clinical trials in sepsis. Intensive Care Med. 2008;34:1955–1960. doi: 10.1007/s00134-008-1274-6

45. Freeman BD, Natanson C. Anti-inflammatory therapies in sepsis and septic shock. Expert Opin Investig Drugs. 2000;9:1651–1663. doi: 10.1517/13543784.9.7.1651

46. R Core Team, R: A Language and Environment for Statistical Computing. R Foundation for Statistical Computing, Vienna, Austria. https://www.R-project.org/ (2024).

47. Balduzzi S, Rücker G, Schwarzer G, How to perform a meta-analysis with R: a practical tutorial, Evidence-Based Mental Health; 22: 153–160. (2019)

